# Detection of clade 2.3.4.4b highly pathogenic H5N1 influenza virus in New York City

**DOI:** 10.1101/2024.04.04.588061

**Authors:** Philip S. Meade, Pooja Bandawane, Kaitlyn Bushfield, Irene Hoxie, Karla R. Azcona, Daneidy Burgos, Sadia Choudhury, Adama Diaby, Mariama Diallo, Kailani Gaynor, Aaron Huang, Kadiatou Kante, Shehryar N. Khan, William Kim, Paul Kehinde Ajayi, Ericka Roubidoux, Sasha Nelson, Rita McMahon, Randy A Albrecht, Florian Krammer, Christine Marizzi

## Abstract

Highly pathogenic avian influenza viruses of the H5N1 clade 2.3.4.4b arrived in North America in the winter of 2021/2022. These viruses have spread across the Americas causing morbidity and mortality in both wild and domestic birds as well as some mammalian species, including cattle. Many surveillance programs in wildlife as well as commercial poultry operations have detected these viruses. Here we conducted surveillance of avian species in the urban environment in New York City. We detected highly pathogenic H5N1 viruses in six samples from four different bird species and performed full genome sequencing. Sequence analysis showed the presence of multiple different genotypes. Our work highlights that the interface between animals and humans that may give rise to zoonotic infections or even pandemics is not limited to rural environments and commercial poultry operations but extends into the heart of our urban centers.

**Importance:** While surveillance for avian influenza viruses is often focused on migratory routes and their associated stop-over locations, or commercial poultry operations, many bird species – including migratory birds – frequent or live in urban green spaces and wetlands. This brings them into contact with a highly dense population of humans and pets providing an extensive urban animal-human interface in which the general public may have little awareness of circulating infectious diseases. This study focuses on virus surveillance at this interface, combined with culturally responsive science education and community outreach.

## Introduction

Zoonotic infections with highly pathogenic avian influenza (HPAI) viruses of the H5N1 subtype were first detected in Hong Kong in 1997 (1, 2). After a hiatus, human infections with these A/goose/Guangdong/1/96-like viruses returned in 2003 (3). Their range was initially restricted to birds in Southeast Asia, but they spread west into the Middle East (4, 5), Europe (6–8) and Africa (8, 9) via migratory birds. In addition, these H5N1 viruses also diversified and split into many different lineages. Between 2010 and 2011 clade 2.3.4.4 viruses emerged in China and started to reassort with other avian influenza viruses producing H5NX genotypes of which many seemed to be of lower pathogenicity (10). These viruses were introduced to the United States (US) in 2014 and caused widespread issues in the poultry industry (e.g. (10)). However, in 2015 they disappeared from circulation in North America (11). A subclade of clade 2.3.4.4, namely clade 2.3.4.4b, spread in Eurasia and Africa in 2020, this time again with an N1 neuraminidase (NA) (12), and arrived in North America via migratory birds in the winter of 2021/2022 (13–15). The clade 2.3.4.4b viruses have now spread across the Americas and have heavily impacted wild bird populations and have hurt the poultry industry (16–19). In addition, infections in mammals – often leading to neurological symptoms and fatal outcomes – have been reported. This includes predatory animals and scavengers feeding on sick or dead birds (20–22). These are mostly seen as dead-end hosts. Marine mammals have also been affected, especially in South America, and mammal-to-mammal transmission is suspected in some of these outbreaks (23–25). Furthermore, clade 2.3.4.4b H5N1 seems to have caused outbreaks in fur farms in Europe in mink and foxes with potential mammal-to-mammal transmission (26–28) and recently reported cases in dairy cattle are also raising concerns. Human cases with clade 2.3.4.4b H5N1 so far have been rare, and only two severe infections are known in the Americas (with a low number of additional ones in Asia including fatalities), which is remarkable given the extent of the spread of this virus and the potential exposure of humans (29–31).

Nevertheless, it is very important to track the spread of this virus to determine potential risk to humans. There is a need for viral surveillance in urban areas which often provide plenty of green space and wetlands for both resident and migratory birds. This, in combination with high human population densities, creates an extensive urban animal-human interface. In this interface, pets can also be impacted as shown by infections of cats and dogs by H5N1 (32–35). Communicating this risk to urban populations is critical. Here, we set out to detect HPAI H5N1 viruses in New York City using surveillance in wildlife rehabilitation centers as well as sampling bird feces from the environment. Our approach is based on collaboration between research institutions, a science outreach organization, wildlife welfare non-profit organizations and community scientists. Community scientists working with our research team have previously reported the first detection of avian paramyxovirus 1 in New York City’s pigeon population (36). The growing interest in biodiversity protection and citizen science has resulted in initiatives that collect a massive quantity of data about birds (37, 38). However, these approaches are frequently limited to participatory data collection (38, 39). In the collaborative New York City Virus Hunters initiative described here, we aim to engage the community in every step of the research process. A core element is mentored research for high school students that self-identify as members of racial or ethnic minoritized groups in science. The students work alongside expert mentors and actively engage in overall study design before safely participating in sample collection, processing, data analysis, dissemination of results and community outreach. The outputs of this program benefit all participants (40).

## Results

### Surveillance strategy and virus detection

Our sampling strategy prioritized samples collected from birds known to contract HPAI H5N1 - principally wild aquatic avian species of *Anseriformes* (ducks, geese and swans), *Charadriiformes* (gulls, terns, auks and other shorebirds) and raptors, such as *Accipitriformes* (hawks, ospreys and other birds of prey) and *Falconidae* (falcons, kestrels). Samples for this study were collected from January 2022 to November 2023. In total 1927 samples were collected and processed for this study. We used two sampling streams. First, 125 environmental fecal samples were collected from New York City parks and green spaces using proper personal protective equipment (N95 masks and gloves). In addition, professional animal rehabilitators at the Wild Bird Fund (WBF) and veterinarians of the Animal Care Centers (ACC) of New York City provided four water samples (3 ml each) and 1798 cloacal (CS), oropharyngeal (OS), and fecal swabs from urban wild and domestic birds submitted to them. From these 1798 samples (237 fecal samples, 783 CS and 764 OS samples, and 14 samples where CS or OP was non-specified; from 895 birds), six were found positive for HPAI H5N1. While for environmental fecal samples collected in urban parks and green spaces the avian species is hard to determine by appearance of the sample, CS, OS, and fecal samples provided by wildlife rehabilitation centers were documented to be from 80 different species (see **Table 1**). The majority were from gulls and terns (348 samples/19.35%), chicken (306 samples/17.01%), geese (247 samples/14.29%), ducks (133 samples/7.39%), hawks (112 samples/6.22%), crows and ravens (85 samples/4.72%), falcons and kestrels (53 samples/2.94%) and cormorants (43 samples/2.39%). Eighty-nine samples (4.94%) were from non-specified species. The remaining samples and the species they were collected from are also listed in **Table 1**. RNA was extracted from 1927 samples and reverse transcription was used to generate cDNA. We then screened the cDNA preparations via a multiplex PCR using primers for the matrix (M) genomic segment, the nucleoprotein (NP) genomic segment and for the hemagglutinin (HA) genomic segment. Primers for the HA segment were H5 HA specific while M and NP primers were designed to detect all known influenza A viruses. Gene products were sent for Sanger sequencing. If Sanger sequencing indicated the presence of influenza A virus, full genome sequencing was performed. Samples from six birds were found to be positive for HPAI H5N1, and full genomes could be sequenced. No environmental fecal samples were found positive for HPAI H5N1. No other avian influenza viruses were detected.

**Table 1:** Details on samples for virological analysis collected from different avian species present at Wild Bird fund or Animal Care Centers of New York City.

|  | Host Species | Scientific name | Order Taxa | Family Taxa | Total samples | HPAI H5N1 positive samples |
| --- | --- | --- | --- | --- | --- | --- |
| 1 | American bittern | *Botaurus lentiginosus* | Pelecaniformes | Ardeidae | 2 |  |
| 2 | Coopers hawk | *Accipiter cooperii* | Accipitriformes | Accipitridae | 22 |  |
| 3 | Hawk | *Accipitridae sp.* | Accipitriformes | Accipitridae | 2 |  |
| 4 | Red-shouldered hawk | *Buteo lineatus* | Accipitriformes | Accipitridae | 4 |  |
| 5 | Red-tailed hawk | *Buteo jamaicensis* | Accipitriformes | Accipitridae | 81 | 1 |
| 6 | Sharp-shinned hawk | *Accipiter striatus* | Accipitriformes | Accipitridae | 3 |  |
| 7 | Dovekie | *Alle alle* | Charadriiformes | Alcidae | 2 |  |
| 8 | American black duck | *Anas rubripes* | Anseriformes | Anatidae | 2 |  |
| 9 | Black scoter | *Melanitta americana* | Anseriformes | Anatidae | 2 |  |
| 10 | Brandt goose | *Branta bernicla* | Anseriformes | Anatidae | 12 |  |
| 11 | Canada goose | *Branta canadensis* | Anseriformes | Anatidae | 222 | 3 |
| 12 | Domestic duck | *Anas p. domesticus* | Anseriformes | Anatidae | 8 |  |
| 13 | Domestic goose | *Anas a. domesticus* | Anseriformes | Anatidae | 2 |  |
| 14 | Duck | *Anas sp.* | Anseriformes | Anatidae | 98 |  |
| 15 | Goose | *Anatidae sp.* | Anseriformes | Anatidae | 10 |  |
| 16 | Mallard | *Anas platyrhynchos* | Anseriformes | Anatidae | 92 |  |
| 17 | Muscovy duck | *Cairina moschata* | Anseriformes | Anatidae | 3 |  |
| 18 | Mute swan | *Cygnus olor* | Anseriformes | Anatidae | 43 |  |
| 19 | Northern shoveler | *Spatula clypeata* | Anseriformes | Anatidae | 3 |  |
| 20 | Ruddy Duck | *Oxyura jamaicensis* | Anseriformes | Anatidae | 9 |  |
| 21 | Snow goose | *Anser caerulescens* | Anseriformes | Anatidae | 11 |  |
| 22 | Swan | *Cygnus sp.* | Anseriformes | Anatidae | 5 |  |
| 23 | Wood duck | *Aix sponsa* | Anseriformes | Anatidae | 13 |  |
| 24 | Great blue heron | *Ardea herodias* | Pelecaniformes | Ardeidae | 6 |  |
| 25 | Green heron | *Butorides virescens* | Pelecaniformes | Ardeidae | 1 |  |
| 26 | Least bittern | *Ixobrychus exilis* | Pelecaniformes | Ardeidae | 2 |  |
| 27 | Night Heron | *Ardeidae sp.* | Pelecaniformes | Ardeidae | 4 |  |
| 28 | Yellow-crowned night heron | *Nyctanassa violacea* | Pelecaniformes | Ardeidae | 8 |  |
| 29 | Homing pigeon | *Columba livia domestica* | Columbiformes | Columbidae | 1 |  |
| 30 | Mourning Dove | *Zenaida macroura* | Columbiformes | Columbidae | 7 |  |
| 31 | Rock pigeon | *Columba livia* | Columbiformes | Columbidae | 41 |  |
| 32 | American crow | *Corvus brachyrhynchos* | Passeriformes | Corvidae | 51 |  |
| 33 | Common raven | *Corvus corax* | Passeriformes | Corvidae | 8 |  |
| 34 | Crow | *Corvus sp.* | Passeriformes | Corvidae | 2 |  |
| 35 | Fish crow | *Corvus ossifragus* | Passeriformes | Corvidae | 18 |  |
| 36 | Raven | *Corvus sp.* | Passeriformes | Corvidae | 6 |  |
| 37 | American kestrel | *Falco sparverius* | Falconiformes | Falconidae | 39 |  |
| 38 | Kestrel | *Falco sp.* | Falconiformes | Falconidae | 2 |  |
| 39 | Peregrine Falcon | *Falco peregrinus* | Falconiformes | Falconidae | 12 | 1 |
| 40 | Loon | *Gavia sp.* | Gaviiformes | Gaviidae | 2 |  |
| 41 | Red-throated loon | *Gavia stellata* | Gaviiformes | Gaviidae | 5 |  |
| 42 | Arctic tern | *Sterna paradisaea* | Charadriiformes | Laridae | 2 |  |
| 43 | Great black-backed gull | *Larus marinus* | Charadriiformes | Laridae | 47 |  |
| 44 | Gull | *Laridae sp.* | Charadriiformes | Laridae | 2 |  |
| 45 | Herring gull | *Larus argentatus* | Charadriiformes | Laridae | 180 |  |
| 46 | Laughing gull | *Leucophaeus atricilla* | Charadriiformes | Laridae | 52 |  |
| 47 | Ring-billed gull | *Larus delawarensis* | Charadriiformes | Laridae | 62 |  |
| 48 | Seagull | *Laridae sp.* | Charadriiformes | Laridae | 3 |  |
| 49 | Guineafowl | *Numida sp.* | Galliformes | Numididae | 2 |  |
| 50 | Osprey | *Pandion haliaetus* | Accipitriformes | Pandionidae | 5 |  |
| 51 | Tufted titmouse | *Baeolophus bicolor* | Passeriformes | Paridae | 1 |  |
| 52 | Northern waterthrush | *Parkesia noveboracensis* | Passeriformes | Parulidae | 1 |  |
| 53 | White-throated sparrow | *Zonotrichia albicollis* | Passeriformes | Passerellidae | 2 |  |
| 54 | House sparrow | *Passer domesticus* | Passeriformes | Passeridae | 6 |  |
| 55 | Cormorant | *Phalacrocorax sp.* | Phalacrocorax | Phalacrocoracidae | 10 |  |
| 56 | Double-crested cormorant | *Nannopterum auritum* | Suliformes | Phalacrocoracidae | 33 |  |
| 57 | Barred rock chicken | *Gallus gallus domesticus* | Galliformes | Phasianidae | 4 |  |
| 58 | Chicken | *Gallus gallus domesticus* | Galliformes | Phasianidae | 302 | 1 |
| 59 | Fowl | *Gallus sp.* | Galliformes | Phasianidae | 36 |  |
| 60 | Japanese quail | *Coturnix japonica* | Galliformes | Phasianidae | 3 |  |
| 61 | Partridge | *Arborophila sp.* | Galliformes | Phasianidae | 12 |  |
| 62 | Pheasant | *Phasianus colchicus* | Galliformes | Phasianidae | 5 |  |
| 63 | Quail | *Coturnix coturnix* | Galliformes | Phasianidae | 11 |  |
| 64 | Turkey | *Phasianidae sp.* | Galliformes | Phasianidae | 4 |  |
| 65 | Wild turkey | *Meleagris gallopavo* | Galliformes | Phasianidae | 7 |  |
| 66 | Yellow-bellied Sapsucker | *Sphyrapicus varius* | Piciformes | Picidae | 4 |  |
| 67 | Grebe | *Podiceps sp.* | Podicipediformes | Podicipedidae | 3 |  |
| 68 | Cory's shearwater | *Calonectris borealis* | Procellariiformes | Procellariidae | 1 |  |
| 69 | Great shearwater | *Ardenna gravis* | Procellariiformes | Procellariidae | 2 |  |
| 70 | Parrot | *Psittacidae sp.* | Psittaciformes | Psittacidae | 3 |  |
| 71 | American coot | *Fulica americana* | Gruiformes | Rallidae | 5 |  |
| 72 | American woodcock | *Scolopax minor* | Charadriiformes | Scolopacidae | 2 |  |
| 73 | Barred owl | *Strix varia* | Strigiformes | Strigidae | 3 |  |
| 74 | Great horned owl | *Bubo virginianus* | Strigiformes | Strigidae | 6 |  |
| 75 | Northern saw-whet owl | *Aegolius acadicus* | Strigiformes | Strigidae | 8 |  |
| 76 | European starling | *Sturnus vulgaris* | Passeriformes | Sturnidae | 1 |  |
| 77 | Gannet | *Sulidae sp.* | Suliformes | Sulidae | 2 |  |
| 78 | Northern Gannet | *Morus bassanus* | Suliformes | Sulidae | 4 |  |
| 79 | American robin | *Turdus migratorius* | Passeriformes | Turdidae | 4 |  |
| 80 | Robin | *Turdidae sp.* | Passeriformes | Turdidae | 3 |  |
|  | Unknown |  |  |  | 89 |  |
|  | Total bird samples |  |  |  | 1798 | 6 |

Of these positive samples, the first (A/Canada goose/New York/NYCVH 22-6038/2022 (H5N1)) was collected from a Canada goose (*Branta canadensis*). This animal was initially found in Hutchinson River Parkway, in the Bronx, and died before the intake exam in August 2022. The next positive sample (A/red-tailed hawk/New York/NYCVH 22-8477/2022 (H5N1))) was derived in October 2022 from a red-tailed hawk (*Buteo jamaicensis*) that was found in close proximity to a major highway in Queens. The bird displayed neurological symptoms at the clinic. In December 2022 two birds found in Brooklyn tested positive for HPAI H5N1. One Canada goose (A/Canada goose/New York/NYCVH 22-9190/2022 (H5N1))), that displayed neurological symptoms and cloudy eyes, and one peregrine falcon (*Falco peregrinus*, strain name A/peregrine falcon/New York/NYCVH 160820/2022). The fifth sample (strain name: A/Canada goose/New York/NYCVH 23-453/2023 (H5N1))) came from a Canada goose found in February of 2023 in Queens. The sixth positive sample (A/chicken/New York/NYCVH 168127/2023 (H5N1))) was collected in April 2023 from a chicken (*Gallus gallus domesticus*) that was found in upper Manhattan (**Figure 1 and Table 2**). No additional positive samples/birds were detected from April 2023 to November 2023. To further analyze our detected viruses, we performed a multiple sequence alignment of their amino acid sequences, and mapped amino acid (AA) changes from the HPAI H5N1 strain A/bald eagle/FL/W22-134-OP/2022 (accession number UWI70064) (41) onto an HA structure from A/chicken/Vietnam/4/2003 (42) (**Figure 3**). The detected AA differences mainly fell outside receptor-binding site and antigenic sites of H5N1 (43, 44), except for T71I. Most differences were only found in one of our NYCVH strains, except for T71I, which was present in all NYCVH strains. It should be noted that isoleucine (I) was present at this position in all 50 strains used to construct our phylogenetic tree, and it is atypical for A/bald eagle/FL/W22-134-OP/2022 to have a threonine (T) at this position. To our knowledge, none of the amino acid changes relative to A/bald eagle/FL/W22-134-OP/2022 have specifically been implicated with increases in pathogenicity or mammalian adaptation.

**Figure 1.**
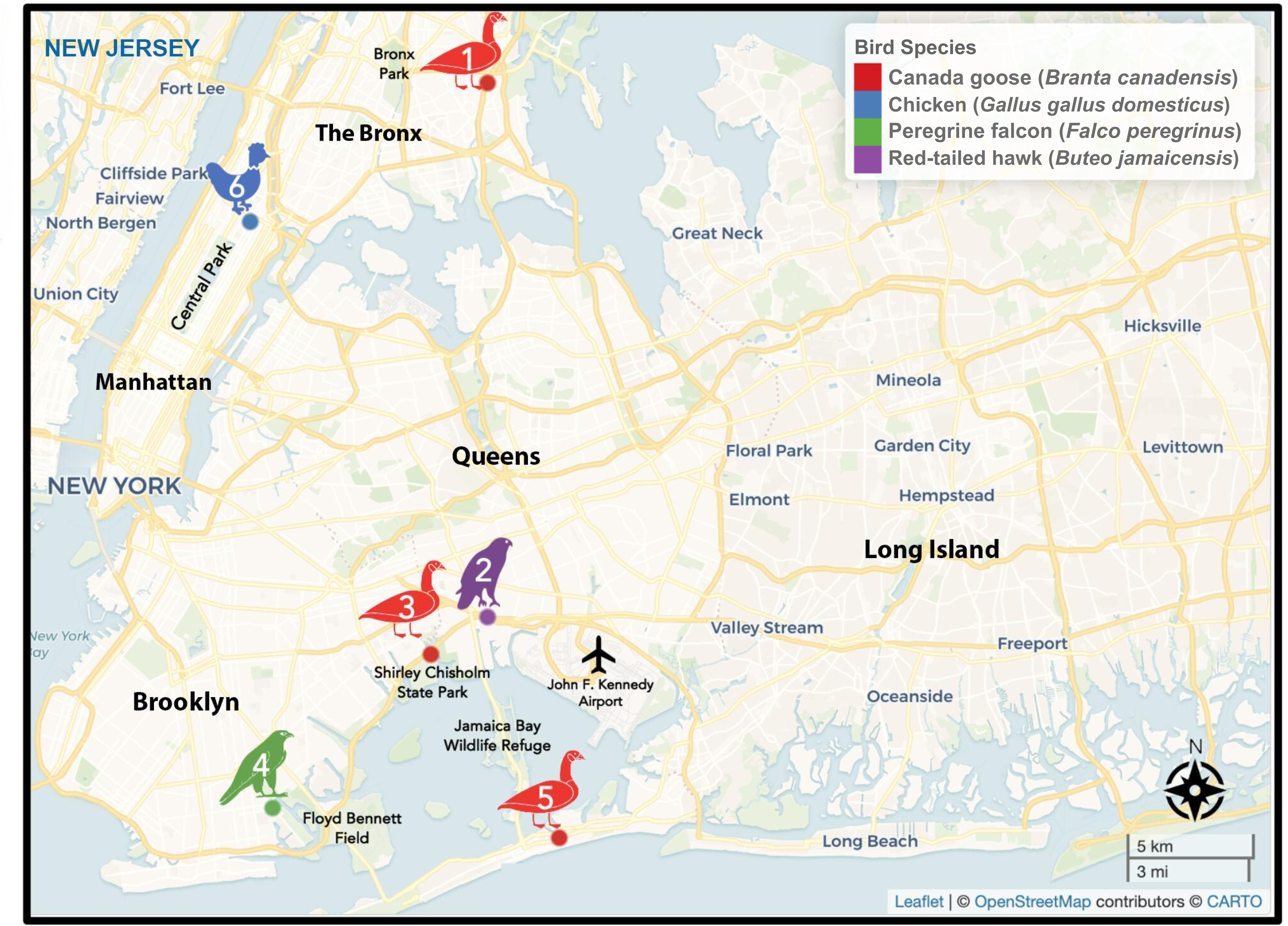
Sampling location of birds that confirmed positive for highly pathogenic avian influenza H5N1 virus (HPAI H5N1) in New York City. This map illustrates the sampling locations of birds that tested positive for the highly pathogenic avian influenza H5N1 virus (HPAI H5N1) in New York City. The approximate locations are plotted based on geocoded addresses (latitude and longitude), providing a visual representation of affected areas. Major parks and natural areas are highlighted in green and labeled for context. The map was created using the leaflet package for mapping visualizations, with additional spatial data handling and aesthetic enhancements performed using the sf, ggplot2, and dplyr packages in RStudio/Posit (Version 2023.09.1+494). The basemap was provided by CARTO, with data sourced from OpenStreetMap under the Open Data Commons Open Database License (ODbL) by the OpenStreetMap Foundation (OSMF).

**Table 2.** Clinical and sampling information for wild birds positive on RT-PCR for highly pathogenic avian influenza (H5N1) in New York City from January 2022 to November 2023.

| Sample ID | Sample type | Species (common name) | Species (scientific name) | Location | Sampling date | Clinical signs |
| --- | --- | --- | --- | --- | --- | --- |
| 22-6038 | Cloacal swab | Canada goose | *Branta canadensis* | Corner of Hutchinson River Pkwy. East and Wilkinson Avenue, Bronx, NY | August 24, 2022 | Died before intake exam. |
| 22-8477 | Fecal | Red tailed hawk | *Buteo jamaicensis* | Belt Pkwy/Nassau Expressway, Queens, NY | October 22, 2022 | Neurologic symptoms, loss of leg function, torticollis, glottis wide open |
| 22-9190 | Oropharyngeal swab | Canada goose | *Branta canadensis* | Fountain Avenue, Brooklyn, NY | December 3, 2022 | Oculi uterque, eyes cloudy and occluded, head tremors, ataxia, unable to stand. |
| 160820 | Oropharyngeal swab | Peregrine falcon | *Falco peregrinus* | Corner of Gerritsen Avenue and Avenue V, Brooklyn, NY | December 18, 2022 | unavailable |
| 23-0453 | Oropharyngeal swab | Canada goose | *Branta canadensis* | Beach 73<sup>rd</sup> Street Avene, Queens, NY | February 1, 2023 | Neurologic symptoms, ataxia, severe head tremors, partial torticollis, labored breathing w/ OU cloudy. |
| 168127 | Fecal | Chicken | *Gallus gallus domesticus* | Corner of 120<sup>th</sup> Street and 5th Avenue, Manhattan, NY | April 2, 2023 | unavailable |

Upon confirmation, detections were reported to the United States Department of Agriculture (USDA) and the associated original samples were transferred to Mount Sinai’s BSL3+ select agent facility (Emerging Pathogens Facility (EPF)/BSL-3 Biocontainment CoRE) for storage. Results were also discussed with the New York City Department of Health and Mental Hygiene as well as the Wild Bird Fund and the Animal Care Centers of New York City, following a previously developed internal and external communication strategy (45, 46). Briefly, the strategy aimed to ensure prompt and informed decisions and that all participants, collaborators and stakeholders are kept fully informed. Successful communication of science and public health messages is complex, and it remains an important challenge to reach potentially vulnerable audiences. Our communication aimed to calm potential anxiety by providing information and instill confidence and trust by addressing all questions to the best of our abilities. It has been noted that communications around emerging infectious disease can be improved when it comes from individuals inside the same community as those receiving the information, simply for the fact that they often share the same language, values and beliefs (47). Therefore, it is incredibly important to ensure researchers involved in pandemic preparedness are committed to bidirectional communication, listening and serving the needs of the community. To reach the scientific community and general public alike, involved students shared their results in multiple languages and through multiple channels. These range from live virtual events and talks at community boards, to in-person symposia and presentations at scientific conferences. Results were also presented to the public at three student research symposia, including the New York City Virus Hunters Symposium on May 31^st^, 2023.

### Phylogenetic analysis of detected HPAI H5N1 genomes

Phylogenetic analysis of the six viral genomes and genotype assignment was performed. The H5 and N1 genes of all six viruses were all typical of the currently circulating 2.3.4.4b clade in the Americas. HA sequences of A/Canada goose/New York/NYCVH 22-6038/2022, A/red-tailed hawk/New York/NYCVH 22-8477/2022, A/Canada goose/New York/NYCVH 22-9190/2022 and A/peregrine falcon/New York/NYCVH 160820/2022 clustered closely together in a tree constructed of 50 HA sequences randomly selected from a list of all available H5N1 strains collected since January 1^st^ 2020 on NCBI’s influenza virus database, downloaded on October 26^th^ 2023 (**Figure 2**). They also cluster with contemporary H5 sequences from 2022 from Ohio, North Carolina but also Colombia. Similarly, their NA sequences cluster together next to the NA sequences of the Ohio and Colombia isolates for which the HAs cluster as well. The two 2023 sequences A/chicken/New York/NYCVH 168127/2023 and A/Canada goose/New York/NYCVH 23-453/2023 are clustering together as well and form their own branch close to a cluster of sequences from 2022 and 2023 North and South American isolates. The NA sequences of these two viruses cluster together, but are also located closely to many different isolates from both North and South America.

**Figure 2.**
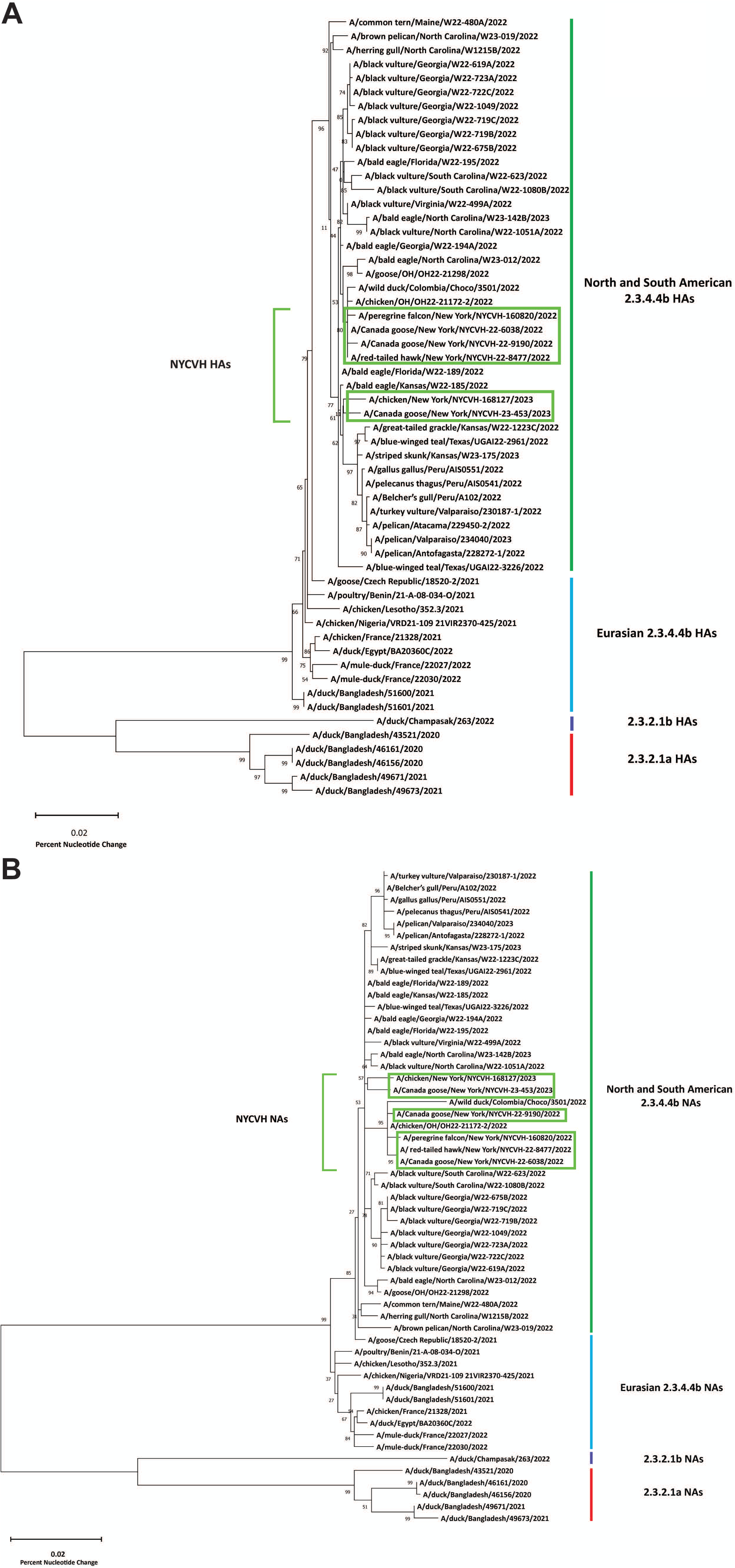
Phylogenetic tree of HA and NA genes of the detected viruses in comparison to sequences from GenBank. HA (**A**) and NA (**B**) gene sequences from 50 strains of H5N1 influenza virus were randomly selected from all available strains collected between January 1 2020 and October 26 2023 (NCBI influenza virus database, https://www.ncbi.nlm.nih.gov/genomes/FLU/Database/nph-select.cgi). The 50 randomly selected H5N1 sequences and the 6 NYCVH H5N1 sequences were used to create a phylogenetic tree in MEGA 11 (https://www.megasoftware.net) using the maximum likelihood method and Tamura-Nei model (65) with a bootstrap test (n=100). Bootstrap support values are shown by branch nodes. Scale bar indicates percent difference in nucleotide sequence.

**Figure 3.**
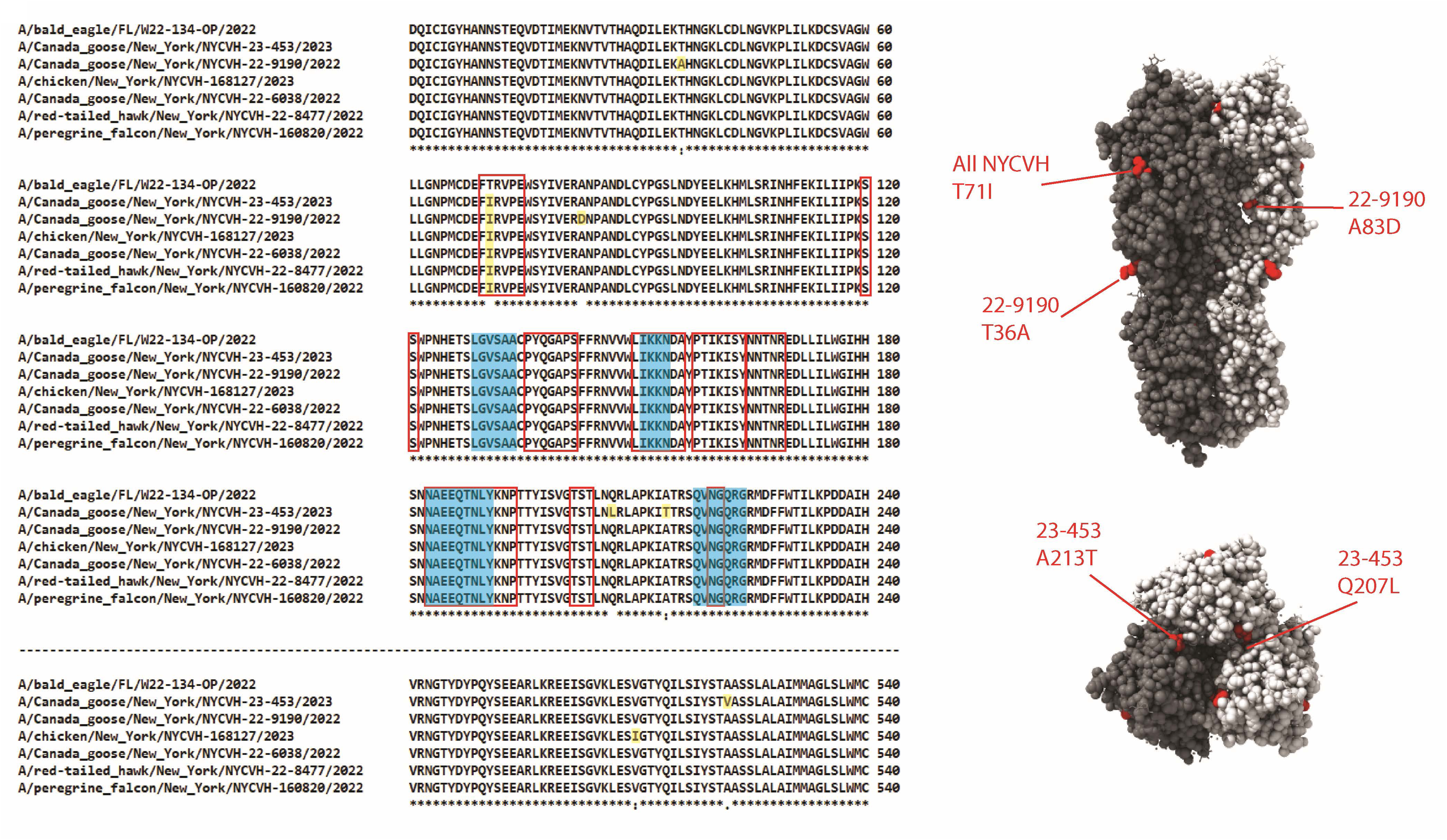
Amino acid sequence analysis of detected HPAI H5N1 strains. A multiple sequence alignment was performed using the HA sequences of NYCVH-detected strains and A/bald eagle/FL/W22-134-OP/2022 in Clustal Omega. Only areas of the alignment with amino acid differences are shown. Residues which were different in NYCVH strains compared to A/bald eagle/FL/W22-134-OP/2022 were highlighted in red on an H5N1 structure based on A/chicken/Vietnam/4/2003 (PDB #6VMZ) (42), visualized with UCSF ChimeraX. V510I and A522V are changes to internal positions in the head domain. The receptor binding site and antigenic sites are indicated by blue highlighting and red outline, respectively.

To identify the genotypes of internal genes, we used a script provided by Youk *et al.* (16) that allows for classification of segments into lineages and determines a genotype based on the genomic segment composition of a virus. We compared our virus sequences with available full length genome sequences from the New York State, New Jersey and Connecticut area surrounding New York City where many infections were detected (**Supplemental Figure 1**). All detected viruses were re-assortant viruses between the Eurasian (EA) and American (AM) lineages. All HA and NA segments were of course from the EA lineage but segments encoding for internal proteins differed. A/Canada goose/New York/NYCVH 22-6038/2022, A/red-tailed hawk/New York/NYCVH 22-8477/2022, A/Canada goose/New York/NYCVH 22-9190/2022 and A/peregrine falcon/New York/NYCVH 160820/2022 were all determined to be genotype B1.3 with AM lineage polymerase and NP segments, and all other segments from the EA lineage (**Table 3**). B1.3 lineage viruses were also found in New York State and neighboring states (New Jersey, Connecticut) during our observation period.(**Table 4**). The more recent A/chicken/New York/NYCVH 168127/2023 and A/Canada goose/New York/NYCVH 23-453/2023 viruses belonged to lineage B3.3, a lineage also detected in New York State in a turkey vulture in April of 2023..This lineage features PB2, PB1, NP, and NS segments from the AM lineage while, PA, HA, NA and M segments are derived from the EA lineage.

**Table 3:** Genotypes of the detected H5N1 strains. American lineages are abbreviated as “am”, and Eurasian lineages are abbreviated as “ea”. Influenza virus gene segments are abbreviated as follows: polymerase basic 2 (PB2), polymerase basic 1 (PB1), polymerase acidic (PA), hemagglutinin (HA), nucleoprotein (NP), neuraminidase (NA), matrix (M), and nonstructural protein (NS).

| Strain | Genotype | PB2 | PB1 | PA | HA | NP | NA | M | NS |
| --- | --- | --- | --- | --- | --- | --- | --- | --- | --- |
| A/red-tailed hawk/New York/NYCVH 22-8477/2022 | **B1.3** | am1.3 | am1.3 | am1.2 | ea1 | am1.2 | ea1 | ea1 | ea1 |
| A/Canada goose/New York/NYCVH 22-6038/2022 | **B1.3** | am1.3 | am1.3 | am1.2 | ea1 | am1.2 | ea1 | ea1 | ea1 |
| A/Canada goose/New York/NYCVH 22-9190/2022 | **B1.3** | am1.3 | am1.3 | am1.2 | ea1 | am1.2 | ea1 | ea1 | ea1 |
| A/peregrine falcon/New York/NYCVH 160820/2022 | **B1.3** | am1.3 | am1.3 | am1.2 | ea1 | am1.2 | ea1 | ea1 | ea1 |
| A/Canada goose/New York/NYCVH 23-453/2023 | **B3.2** | am2.1 | am1.2 | ea1 | ea1 | am1.4.1 | ea1 | ea1 | am1.1 |
| A/chicken/New York/NYCVH 168127/2023 | **B3.3** | am2.2 | am1.4 | ea1 | ea1 | am1.4.1 | ea1 | ea1 | am1.1 |

**Table 4:** Genotypes of strains detected in New York, New Jersey, and Connecticut from August 2022 – April 2023. American lineages are abbreviated as “am”, and Eurasian lineages are abbreviated as “ea”. Influenza virus gene segments are abbreviated as follows: polymerase basic 2 (PB2), polymerase basic 1 (PB1), polymerase acidic (PA), hemagglutinin (HA), nucleoprotein (NP), neuraminidase (NA), matrix (M), and nonstructural protein (NS).

| Collection date | Strain | Genotype | PB2 | PB1 | PA | HA | NP | NA | M | NS |
| --- | --- | --- | --- | --- | --- | --- | --- | --- | --- | --- |
| 10/11/2022 | A/domestic_duck/New_Jersey/22-032412-001-original | **B1.1** | am1.1 | am1.1 | ea1 | ea1 | am1.2 | ea1 | ea1 | ea1 |
| 10/21/2022 | A/African_goose/New_Jersey/22-033679-001-original | **B1.3** | am1.3 | am1.3 | am1.2 | ea1 | am1.2 | ea1 | ea1 | ea1 |
| 10/28/2022 | A/wild_turkey/New_York/22-034864-002-original | **B1.3** | am1.3 | am1.3 | am1.2 | ea1 | am1.2 | ea1 | ea1 | ea1 |
| 11/1/2022 | A/chicken/New_Jersey/22-035003-009-original | **B1.3** | am1.3 | am1.3 | am1.2 | ea1 | am1.2 | ea1 | ea1 | ea1 |
| 11/3/2022 | A/chicken/New_York/22-035475-001-original | **B2.2** | am1.2 | ea1 | ea1 | ea1 | am1.1 | ea1 | ea1 | am1.2 |
| 11/3/2022 | A/chicken/New_York/22-035476-001-original | **B2.2** | am1.2 | ea1 | ea1 | ea1 | am1.1 | ea1 | ea1 | am1.2 |
| 11/7/2022 | A/Muscovy_duck/New_York/22-036304-003-original | **B1.3** | am1.3 | am1.3 | am1.2 | ea1 | am1.2 | ea1 | ea1 | ea1 |
| 11/7/2022 | A/chicken/New_York/22-036304-007-original | **B1.3** | am1.3 | am1.3 | am1.2 | ea1 | am1.2 | ea1 | ea1 | ea1 |
| 2/8/2023 | A/American_crow/New_York/23-004806-001-original | **B3.5** | am3.2 | am1.2 | ea1 | ea1 | am1.4.1 | ea1 | ea1 | am1.1 |
| 2/10/2023 | A/great_horned_owl/New_York/23-005127-001-original | **B1.2** | am1.2 | ea1 | ea1 | ea1 | am1.1 | ea1 | am1 | am1.1 |
| 2/11/2023 | A/red_tailed_hawk/New_York/23-005536-001-original | **B2.2** | am1.2 | ea1 | ea1 | ea1 | am1.1 | ea1 | ea1 | am1.2 |
| 2/13/2023 | A/American_crow/New_York/23-005128-001-original | **B3.5** | am3.2 | am1.2 | ea1 | ea1 | am1.4.1 | ea1 | ea1 | am1.1 |
| 2/13/2023 | A/peregrine_falcon/New | **B1.3** | am1.3 | am1.3 | am1.2 | ea1 | am1.2 | ea1 | ea1 | ea1 |
|  | _York/23-005700-001-original |  |  |  |  |  |  |  |  |  |
| 2/15/2023 | A/Canada_goose/New_York/23-005698-001-original | **B1.2** | am1.1 | am1.1 | am1 | ea1 | am1.2 | ea1 | ea1 | ea1 |
| 2/15/2023 | A/Canada_goose/New_York/23-005699-001-original | **B1.2** | am1.1 | am1.1 | am1 | ea1 | am1.2 | ea1 | ea1 | ea1 |
| 2/15/2023 | A/Canada_goose/New_York/23-005699-001-original | **B1.2** | am1.1 | am1.1 | am1 | ea1 | am1.2 | ea1 | ea1 | ea1 |
| 2/20/2023 | A/American_crow/New_York/23-005695-001-original | **B3.5** | am3.2 | am1.2 | ea1 | ea1 | am1.4.1 | ea1 | ea1 | am1.1 |
| 2/22/2023 | _A/Canada_goose/New_York/23-006363-001-original | **B3.5** | am3.2 | am1.2 | ea1 | ea1 | am1.4.1 | ea1 | ea1 | am1.1 |
| 2/27/2023 | A/American_crow/New_York/23-006663-001-original | **B3.5** | am3.2 | am1.2 | ea1 | ea1 | am1.4.1 | ea1 | ea1 | am1.1 |
| 2/27/2023 | A/Canada_goose/New_York/23-006664-001-original | **B3.5** | am3.2 | am1.2 | ea1 | ea1 | am1.4.1 | ea1 | ea1 | am1.1 |
| 3/6/2023 | A/Canada_goose/New_York/23-008112-001-original | **B1.2** | am1.1 | am1.1 | am1 | ea1 | am1.2 | ea1 | ea1 | ea1 |
| 3/15/2023 | A/American_crow/New_York/23-009037-001-original | **B3.5** | am3.2 | am1.2 | ea1 | ea1 | am1.4.1 | ea1 | ea1 | am1.1 |
| 3/16/2023 | A/peregrine_falcon/New_York/23-009036-001-original | **B3.5** | am3.2 | am1.2 | ea1 | ea1 | am1.4.1 | ea1 | ea1 | am1.1 |
| 3/17/2023 | A/American_crow/New_York/23-009035-001-original | **B3.5** | am3.2 | am1.2 | ea1 | ea1 | am1.4.1 | ea1 | ea1 | am1.1 |
| 3/23/2023 | _A/red-tailed_hawk/Connecticut/23-009886-001-original | **B1.3** | am1.3 | am1.3 | am1.2 | ea1 | am1.2 | ea1 | ea1 | ea1 |
| 3/23/2023 | A/red-tailed_hawk/Connecticut/23-009880-001-original | **B1.3** | am1.3 | am1.3 | am1.2 | ea1 | am1.2 | ea1 | ea1 | ea1 |
| 4/6/2023 | A/turkey_vulture/New_York/23-011005-001-original | **B3.3** | am2.2 | am1.4 | ea1 | ea1 | am1.4.1 | ea1 | ea1 | am1.1 |
| 4/13/2023 | A/red-shouldered_hawk/New_York/23-012563-001-original | **B3.2** | am2.1 | am1.2 | ea1 | ea1 | am1.4.1 | ea1 | ea1 | am1.1 |
| 4/21/2023 | _A/Canada_goose/New_York/23-013244-001-original | **B1.3** | am1.3 | am1.3 | am1.2 | ea1 | am1.2 | ea1 | ea1 | ea1 |
| 4/27/2023 | _A/American_crow/New_York/23-014010-001-original | **B3.5** | am3.2 | am1.2 | ea1 | ea1 | am1.4.1 | ea1 | ea1 | am1.1 |
| 4/27/2023 | A/Canada_goose/New_York/23-014011-001-original | **B1.3** | am1.3 | am1.3 | am1.2 | ea1 | am1.2 | ea1 | ea1 | ea1 |

## Discussion

The recent spread of the panzootic clade 2.3.4.4b H5N1 across the globe has caused significant damage to wild bird populations and to the poultry industry (16, 41, 48, 49). Spillovers into mammals have caused concerns about mammalian adaptation of this clade. However, despite the wide spread of the clade 2.3.4.4b H5N1 virus and likely significant exposure of humans to it (hunters, poultry farmers etc.), human infections have so far been rare with only two known severe cases in the Americas (29, 30) and a small number in Asia (31). Avian influenza virus surveillance is often carried out in wild birds in rural areas, through hunter programs as well as in domestic poultry operations. However, surveillance systems to detect the virus in urban wild birds is often absent. Despite that, many bird species inhabit or temporarily visit urban areas which in many cases have ample green space as well as aquatic habitats for waterfowl. This is exemplified by the long list of species sampled in this study. Our study focused on this urban space using two sample streams including samples from animal rehabilitation centers (Wild Bird Fund and Animal Care Centers of New York City) and environmental fecal samples sourced via a citizen/community science project (New York City Virus Hunters). Including the community in viral surveillance in a safe way generates interest and understanding of the topic in the population which is important given the science skepticism which has come to light through the coronavirus disease 2019 (COVID-19) pandemic (50, 51).

Our work identified six HPAI H5N1 viruses in 1927samples (corresponding to at least 895 birds). These viruses were found in species known to be susceptible for H5N1 infection. Based on infection patterns in our area, we did expect to find HPAI H5N1 virus in Canada geese (which are highly susceptible to H5N1 infections (52, 53)) as well as in raptors (peregrine falcon, red-tailed hawk) which often get infected when feeding on infected prey or carcasses. While H5N1 is known to infect chickens, it was somewhat unexpected to receive samples from a chicken found in Marcus Garvey Park in Manhattan. Almost all our samples from chickens were from birds in captivity. It remains unclear if this chicken was intentionally released or escaped from captivity elsewhere, as does the context in which it became infected (in captivity or after release). It is important to state that all six positive samples came from either the Wild Bird Fund or the Animal Care Centers of New York City, stressing the important role that urban wildlife rehabilitation centers can play in urban viral surveillance efforts. The detected HA and NA sequences clustered with other H5 and N1 sequences from North and South American clade 2.3.4.4b H5N1 viruses circulating at approximately the same time and they belonged to two different genotypes, which are both reassortants between the Eurasian 2.3.4.4b H5N1 and American avian influenza viruses. It has recently been shown that these reassortants can have increased pathogenicity in mammals as compared to the full Eurasian genotype of 2.3.4.4b H5N1 (41). The genotypes of our NYCVH-detected HPAI H5N1 viruses have also been detected in the region (defined as the states of New York, New Jersey, and Connecticut) during the same time period. Of note, while many infections in mammals have been reported in the Americas including with severe (and often neurological) symptoms and outcomes; most have been ‘dead end’ infections in scavengers or predatory animals which presumably fed on infected birds or bird carcasses (20–22). However, mammal-to-mammal transmission is suspected in several outbreaks in fur farms in Europe and in marine mammals in South America (26–28) and recent cases in cows in several US states and in a goat raise concerns.

Our study shows that clade 2.3.4.4b H5N1 highly pathogenic avian influenza can be present in birds that migrate through or live in urban centers. This highlights the importance of viral surveillance at the urban animal-human interface in which wild animals may potentially interact with a high density population of humans and their pets. Humans may interact with infected birds directly (handling an injured bird) or indirectly (e.g. by coming in contact with feces or contaminated water in parks). Pets including cats and dogs are susceptible to HPAI H5N1 and transmissions from birds to both pet species have happened via contact with infected birds or bird carcasses – scenarios which could occur in urban green spaces where pets are frequently taken (32–35, 54). Our study highlights these risks. However, it needs to be emphasized that a very small number of birds were found positive. Of note, the low percentage of positive animals could also be due to the sensitivity of the screening pipeline used and other assays or tests may produce a higher number of positives.

An important aspect of our work is to involve the population and all stakeholders in surveillance efforts and communicate findings and risks efficiently. In order to do so, we have shared and discussed our results with the New York City Department of Health and Mental Hygiene and we have worked out a communication strategy. Junior Scientists from the New York City Virus Hunters Program have also shared the results of our study with the public during our annual symposium in June of 2023. We believe it is important for the public to understand that HPAI H5N1 may be present in birds, as well as their feces and other secretions in urban spaces, that sick or weirdly behaving birds (or other wildlife) should be reported to the authorities and only be handled by professionals in proper personal protective equipment, and that pets should be kept away from urban wildlife. Furthermore, it is important for physicians in urban centers to know about the potential presence of HPAI H5N1 and be aware that atypical influenza cases in humans may be caused by avian influenza viruses in humans. So far studies suggest that North American clade 2.3.4.4b viruses are susceptible to all classes of influenza drugs which are available as treatment options (41).

In summary, through a science outreach and community science project, we found clade 2.3.4.4b H5N1 highly pathogenic avian influenza viruses in New York City birds. The presence of the virus poses a low but non-zero risk for humans and pets and more awareness about the presence of this virus in the urban animal-human interface is needed.

## Methods and Materials

### Sample collection

No birds were killed for the purposes of this study. Work at Mount Sinai was approved by the Icahn School of Medicine at Mount Sinai Institutional Animal Care and Use Committee (IPROTO202300000038). Sampling in New York City parks was permitted by the New York City Parks and Recreation. WBF operate under a Department of Environmental Conservation license and both WBF and ACC operate under a U.S. Fish & Wildlife Service Migratory Bird permit. Fecal and swab samples from live, sick and recently deceased or euthanized birds were provided by WBF and ACC and were collected by veterinarians or licensed veterinary technicians as part of standard veterinary care of the birds. Water samples (3 ml each) were collected at WBF’s indoor waterfowl rehabilitation pool, using sterile pipettes and stored individually in cryotubes. Sampling focused on aquatic birds particularly *Anseriformes* (including *Anas* ducks, geese and swans), *Ciconiiformes* (including gulls, cormorants and shorebirds) and raptors (including hawks, eagles and falcons). All live bird sampling was performed by New York State licensed wildlife rehabilitators employed by the WBF or ACC. Fecal, oropharyngeal and cloacal swabs were collected from each bird using sterile flocked nylon-tipped swabs and stored individually in cryovials containing either MicroTest viral transport medium (Thermo Scientific, USA) or medium containing 50% phosphate-buffered saline and 50% glycerol, supplemented with 1% Antibiotic-Antimycotic 100X (Gibco, Thermo Scientific, USA). Samples were kept at 4°C for up to 4 hours, stored at -20°C for up to 7 days and then stored at -80°C . Cold chain was maintained throughout delivery of samples to the laboratory. Environmental fecal samples that appeared fresh (still moist) were collected opportunistically in urban parks and greenspaces where birds were observed congregating, sacrificing the specific identity of the birds being sampled. Environmental fecal samples included in this study were collected on the following locations and dates (12 sampling field trips total): Manhattan, New York: Central Park (May 2022, January and November 2023), Riverside Park (April and June 2023),Tompkins Square Park (April 2022 and May 2023) and South Bronx, New York: Saint Mary’s Park (October 2023). For each location samples were obtained over a wide area of interest, and not a single point. To avoid sampling the same bird more than once, samples were collected with a minimum distance of 20 cm between each other. A transect sampling strategy was employed when sampling around bodies of water, like city ponds. All samples were collected and preserved in the same manner as those collected from live birds. When possible, samples were collected avoiding visible uric acid and soil to prevent potential contamination of PCR inhibitors.

### RNA extraction and RT-PCR

Fecal samples were diluted in phosphate-buffered saline, pH 7.4 (1X, Thermo Scientific, USA) for processing. Suspended fecal samples, oropharyngeal swabs and cloacal swabs, were centrifuged at 4,000 × g for 15 min and viral RNA was extracted from each supernatant using the QIAamp Viral RNA minikit (Qiagen, USA) according to the manufacturer’s instructions. The Stool Total RNA purification kit (Norgen Biotek Corporation, Canada) was also used to extract RNA from fecal samples. Samples collected from the same bird were not pooled. Conventional two step reverse transcriptase polymerase chain reaction (RT-PCR) was employed using the Invitrogen SuperScript IV first-strand synthesis system (Thermo Scientific, USA) for cDNA synthesis and DreamTaq Green PCR Master Mix (2X) (Thermo Scientific, USA) for RT-PCR. First, cDNA was synthesized using a minimum of 250 ng of RNA at 55°C for 10 min using a previously described universal primer (Uni12, *AGCAAAAGCAGG* (55)). Then cDNA was amplified using previously described primers for HPAI H5N1 surveillance that target the nucleoprotein (NP, NP1200 Forward, *CAGRTACTGGGCHATAAGRAC* and NP1529 Reverse, *GCATTGTCTCCGAAGAAATAAG* (56)), matrix (M, M52C Forward, *CTTCTAACCGAGGTCGAAACG* and M253R Reverse, *AGGGCATTTTGGACAAAKCGTCTA* (*56*)) and hemagglutinin (H5, H5.2344-1673 Forward, *TACCAAATAYTGTCAATTTATTCAAC* and H5.2344-1749 Reverse, *GTAAYGACCCRTTRGARCACATCC* (*57*)) genes. Primers for HA (H5) were included for prompt identification of HPAI H5N1 viruses, facilitating quick notification of partner organizations as necessary to handle infected birds. Cycling conditions for the multiplex PCR consisted of a pre-denaturation step at 95°C for 1 min, followed by 30 cycles of denaturation at 95°C for 1 min, annealing at 45°C for 30 s, and extension at 72°C for 30 s, with a final extension step at 72°C for 5 min. PCR amplicons were visualized with SYBR Safe DNA Gel Stain in 2% Ultra Pure Agarose (Thermo Scientific, USA). DNA bands were excised and purified using the QIAquick Gel Extraction Kit (Qiagen, USA) and sent for commercial Sanger sequencing (through Genewiz, New Jersey facility) to confirm the identity of samples that screened positive. Samples that screened positive for H5 HA and also were identified as H5 by Sanger sequencing (to exclude false positives) were reported to the USDA. The remaining sample material was moved to Mount Sinai’s select agent facility for storage.

### Next generation sequencing

Samples which tested positive for H5, NP and M by PCR were then used for next generation sequencing. RT-PCR products were quantified on a Qubit 4 Fluorometer using HS DNA reagents (Invitrogen). A volume of 3.5 µL of the cDNA product was used in a 50 µL PCR reaction with Phusion™ High-Fidelity DNA Polymerase (2 U/µL) (ThermoFisher). Three universal Influenza A primers at a 0.20 μM concentration were used in the PCR reaction. Commonuni13 (GCCGGAGCTCTGCAGATATCAGTAGAAACAAGG), Commonuni12G (GCCGGAGCTCTGCAGATATCAGCGAAAGCAGG) and Commonuni12A (GCCGGAGCTCTGCAGATATCAGCA AAAGCAGG), 0.2 μM dNTPs and 1X HF buffer were also components of the reaction. Single reaction multiplex PCR was performed for sample amplification. Amplification occurred under the following cycling parameters: samples were initially denatured at 94°C for 2 minutes, then underwent 5 cycles of 94°C for 30 seconds, 45°C for 30 seconds and 60°C for 3 minutes. Following these cycling parameters 31 cycles of 94°C for thirty seconds, 57°C for 30 seconds, and 68°C for 3 minutes, followed by a 4°C hold. PCR products underwent DNA purification and size selection using AMpure XP beads. Library preparation was performed using the NEXTRA XT DNA Library Preparation Kit (Illumnia, CA, USA) following manufacturer protocol to generate multiplex paired end sequencing libraries. Post fragmentation automated electrophoresis was performed using the Tape Station 1450 (Agilent Technologies). Sample libraries were quantified using a Qubit 4 Fluorometer (Invitrogen). Sample molarity was determined according to the following formula:

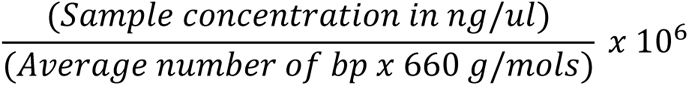

Samples were pooled using Illumina’s pooling calculator in equimolar amounts, creating a paired end fragmented library pool in which each sample is represented with unique indices. Upon pooling and diluting samples to 4nM using 8.5pH 10mM tris(hydroxymethyl)aminomethane (Tris)-HCl with 0.1% Tween20, ‘Protocol A: Standard Normalization method’ from the ‘illlumina MiSeq System: Denature and Dilute Libraries Guide’ was followed. Samples were sequenced using a Miseq device (Illumnia, CA, USA) at a final concentration of 12 pM with a 15% PhiX spike in. After the 2 x 300bp Miseq paired-end sequencing run, the instrument performed base calling on the data, and collected reads with matching indices to generate paired end forward and reverse read Illumina FASTQ files for each sample.

Illumina reads in the form of FASTQ files are produced by the Miseq with adapter sequences on the 5’ and 3’ ends for sample identification, therefore the first step in analysis was to remove the non-viral DNA. FASTQ files were uploaded to the Galaxy web platform, and several tools available through the usegalaxy.org public server (58), were used for genome assembly. Cutadapt (Martin) was used to removed adaptor sequences and to generate FASTA files for analysis. Metagenome *de novo* assembly was performed using the metaSPAdes approach. Nucleotide to protein BLAST (NCBI) was performed on generated sequences to confirm identity and relevance (59). Influenzas A virus segments were identified by NCBI Flu annotator (60). Nucleotide to nucleotide BLAST was run to determine reference sequences. FASTA files were then indexed and mapped to the reference genome using BWA-MEM2 (61) and *Simple Illumina* parameters. BWA-MEM2 produces sorted and indexed BAM files. To account for potential laboratory-derived influenza virus contaminants that could alter the final consensus sequences, we performed the same analysis on a filtered read set, generated by taking the unaligned reads leftover after aligning the data to the A/Puerto Rico/8/1934 reference genome (NCBI Taxonomy ID id183764) using Bowtie2 (62). To ensure only reads definitively derived from known contaminants were excluded, and no reads derived from influenza virus from the sample were discarded, we adjusted the parameters of the alignment algorithm. To increase strictness, the alignment was run in end-to-end mode and increased the mismatch penalties, gap opening and extension penalties, and used the maximum seed substring length allowing for zero sequence mismatches in the seed alignment during multiseed alignment. Final analysis and consensus sequences were produced using the indexed BAM file and Ivar Consensus (63) and Geneious. The antigenic genotypes of the consensuses sequences were confirmed using NCBI Flu annotator. Nucleotide to protein BLAST (NCBI) was performed on generated sequences to confirm identity and relevance. Sequence relevance is determined by the location of isolation and date of isolation in terms of the closest reference sequence, as well as its similarity to existing reference sequences for HPAI H5N1 viruses from 2022 forward.

### Phylogenetic analysis

A phylogenetic analysis comparing the sequences obtained from next generation sequencing with other recent H5N1 sequences was performed. All available HA and NA sequences from H5N1 strains collected since January 1^st^, 2020 were downloaded as FASTA files from the NCBI influenza virus database (https://www.ncbi.nlm.nih.gov/genomes/FLU/Database/nph-select.cgi) on October 26 2023. 50 pairs of HA and NA sequences (from the same influenza virus strain) were randomly selected for inclusion in the phylogenetic tree. In addition to our squencs, the following sequences were used: A/black vulture/Georgia/W22-1049/2022:, WEV84579, WEV84580; A/bald eagle/Florida/W22-189/2022: OP221398, OP221399; A/mule-duck/France/22030/2022: OQ632831, OQ632862; A/duck/Bangladesh/49673/2021: OP023902, OP023904; A/duck/Egypt/BA20360C/2022: OP590397, OP590399; A/duck/Bangladesh/49671/2021: OP023710, OP023712; A/black vulture/North Carolina/W22-1051A/2022: OQ694936, OQ694937; A/goose/Czech Republic/18520-2/2021: OL638145, OL638147; A/chicken/Lesotho/352.3/2021: OL477524, OL477526; A/turkey vulture/Valparaiso/230187-1/2022: OR125345, OR125347; A/black vulture/Georgia/W22-719C/2022: OQ584791, OQ584792; A/striped skunk/Kansas/W23-175/2023: OQ954544, OQ954545; A/duck/Bangladesh/51601/2021: OP030702, OP030704; A/blue-winged teal/Texas/UGAI22-3226/2022: OQ733076, OQ733077; A/blue-winged teal/Texas/UGAI22-2961/2022: OQ733108, OQ733109; A/pelican/Atacama/229450-2/2022: OR125340, OR125342; A/black vulture/Georgia/W22-723A/2022: OQ600260, OQ584498; A/chicken/Nigeria/VRD21-109_21VIR2370-425/2021: MW961460, MW961462; A/black vulture/South Carolina/W22-1080B/2022: OQ694870, OQ694871; A/pelecanus thagus/Peru/AIS0541/2022: OQ547335, OQ547337; A/black vulture/Georgia/W22-675B/2022: OQ584544, OQ584545; A/black vulture/Georgia/W22-619A/2022: OQ584552, OQ584553; A/black vulture/South Carolina/W22-623/2022: OQ584575, OQ584576; A/mule-duck/France/22027/2022: OQ632829, OQ632861; A/chicken/France/21328/2021: OQ632895, OQ632900; A/great-tailed grackle/Kansas/W22-1223C/2022: OQ734910, OQ734911; A/pelican/Antofagasta/228272-1/2022: OR125399, OR125401; A/black vulture/Virginia/W22-499A/2022: OP377388, OP377389; A/pelican/Valparaiso/234040/2023: OR125162, OR125164; A/duck/Bangladesh/43521/2020: MW466215, MW466211; A/black vulture/Georgia/W22-722C/2022: OQ584606, OQ584607; A/wild duck/Colombia/Choco/3501/2022: OQ683498, OQ683500; A/duck/Bangladesh/46156/2020: OM938314, OM938316; A/goose/OH/OH22-21298/2022: OR136609, OR136611; A/duck/Bangladesh/46161/2020: OM938292, OM938294; A/herring gull/North Carolina/W1215B/2022: OQ734937, OQ734938; A/bald eagle/North Carolina/W23-142B/2023: OQ732988, OQ732989; A/brown pelican/North Carolina/W23-019/2022: OQ734918, OQ734919; A/poultry/Benin/21-A-08-034-O/2021: ON870434, ON943071; A/bald eagle/Florida/W22-195/2022: OP221327, OP221328; A/bald eagle/Georgia/W22-194A/2022: OP221382, OP221383; A/duck/Bangladesh/51600/2021: OP030710, OP030712; A/gallus gallus/Peru/AIS0551/2022: OQ547415, OQ547417: A/bald eagle/Kansas/W22-185/2022: OP377646, OP377647; A/bald eagle/North Carolina/W23-012/2022: OQ982396, OQ982397; A/common tern/Maine/W22-480A/2022: OP377502, OP377503; A/black vulture/Georgia/W22-719B/2022: OQ737753, OQ737754; A/chicken/OH/OH22-21172-2/2022: OR136572, OR136574; A/duck/Champasak/263/2022: OR105066, OR105068; A/Belcher’s_gull/Peru/A102/2022: OQ747759, OQ747766. The sequences were aligned with MUSCLE(64), and were manually trimmed to remove non-coding regions before and after the protein sequence. The tree was created in MEGA 11 (https://www.megasoftware.net) using the Maximum Likelihood method (65) with a bootstrap test (n=100). Initial trees were generated with the Neighbor-Join and BioNJ algorithms applied to a matrix of pairwise distances created using the Tamura-Nei model, and the topology with the superior log likelihood value was selected. Multiple sequence alignment of amino acid sequences was performed with Clustal Omega v1.2.4. Protein structure was visualized with UCSF ChimeraX, using the publicly available H5 structure #6V (42).

### Genotyping and survey of HPAI H5N1 detected in New York, New Jersey, and Connecticut

Genotyping was carried out according to a method and data pipeline established by Youk *et al.*, (16) (https://github.com/USDA-VS/GenoFLU). HPAI H5N1 sequences from samples collected in New York, Connecticut, and New Jersey between August 2022 and April 2023 were downloaded on March 26, 2024 from Global Initiative for Sharing All Influenza Database (GISAID). HPAI H5N1 detection data for August 2022 – April 2023 was downloaded on March 24, 2024 from the United States Department of Agriculture, Animal and Plant Health Inspection Service (USDA APHIS) database on wild bird HPAI H5N1 detections. The following sequences were used: A/domestic_duck/New_Jersey/22-032412-001-original: EPI2264144, EPI2264145, EPI2264143, EPI2264147, EPI2264140, EPI2264146, EPI2264142, EPI2264141; A/African_goose/New_Jersey/22-033679-001-original: EPI2263775, EPI2263776, EPI2263774, EPI2263778, EPI2263771, EPI2263777, EPI2263773, EPI2263772; A/wild_turkey/New_York/22-034864-002-original: EPI2263599, EPI2263600, EPI2263598, EPI2263602, EPI2263595, EPI2263601, EPI2263597, EPI2263596; A/chicken/New_Jersey/22-035003-009-original: EPI2263463, EPI2263464, EPI2263462, EPI2263466, EPI2263459, EPI2263465, EPI2263461, EPI2263460; A/chicken/New_York/22-035475-001-original: EPI2260707, EPI2260708, EPI2260706, EPI2260710, EPI2260703, EPI2260709, EPI2260705, EPI2260704; A/chicken/New_York/22-035476-001-original: EPI2260715, EPI2260716, EPI2260714, EPI2260718, EPI2260711, EPI2260717, EPI2260713, EPI2260712; A/Muscovy_duck/New_York/22-036304-003-original: EPI2260795, EPI2260796, EPI2260794, EPI2260798, EPI2260791, EPI2260797, EPI2260793, EPI2260792; A/chicken/New_York/22-036304-007-original: EPI2260803, EPI2260804, EPI2260802, EPI2260806, EPI2260799, EPI2260805, EPI2260801, EPI2260800; A/American_crow/New_York/23-004806-001-original: EPI2613579, EPI2613580, EPI2613578, EPI2613582, EPI2613575, EPI261358, EPI2613577, EPI2613576; A/great_horned_owl/New_York/23-005127-001-original: EPI2613603, EPI2613604, EPI2613602, EPI2613606, EPI2613599, EPI2613605, EPI2613601, EPI2613600; A/red-tailed_hawk/New_York/23-005536-001-original: EPI2613635, EPI2613636, EPI2613634, EPI2613638, EPI2613631, EPI2613637, EPI2613633, EPI2613632; A/American_crow/New_York/23-005128-001-original: EPI2613611, EPI2613612, EPI2613610, EPI2613614, EPI2613607, EPI2613613, EPI2613609, EPI2613608; A/peregrine_falcon/New_York/23-005700-001-original: EPI2613699, EPI2613700, EPI2613698, EPI2613702, EPI2613695, EPI2613701, EPI2613697, EPI2613696; A/Canada_goose/New_York/23-005698-001-original: EPI2613683, EPI2613684, EPI2613682, EPI2613686, EPI2613679, EPI2613685, EPI2613681, EPI2613680; A/Canada_goose/New_York/23-005699-001-original: EPI2613691, EPI2613692, EPI2613690, EPI2613694, EPI2613687, EPI2613693, EPI2613689, EPI2613688; A/American_crow/New_York/23-005695-001-original: EPI2613675, EPI2613676, EPI2613674, EPI2613678, EPI2613671, EPI2613677, EPI2613673, EPI2613672; A/Canada_goose/New_York/23-006363-001-original: EPI2613779, EPI2613780, EPI2613778, EPI2613782, EPI2613775, EPI2613781, EPI2613777, EPI2613776; A/American_crow/New_York/23-006663-001-original: EPI2613819, EPI2613820, EPI2613818, EPI2613822, EPI2613815, EPI2613821, EPI2613817, EPI2613816; A/Canada_goose/New_York/23-006664-001-original: EPI2613827, EPI2613828, EPI2613826, EPI2613830, EPI2613823, EPI2613829, EPI2613825, EPI2613824; A/Canada_goose/New_York/23-008112-001-original: EPI2614003, EPI2614004, EPI2614002, EPI2614006, EPI2613999, EPI2614005, EPI2614001, EPI2614000; A/American_crow/New_York/23-009037-001-original: EPI2613979, EPI2613980, EPI2613978, EPI2613982, EPI2613975, EPI2613981, EPI2613977, EPI2613976; A/peregrine_falcon/New_York/23-009036-001-original: EPI2613971, EPI2613972, EPI2613970, EPI2613974, EPI2613967, EPI2613973, EPI2613969, EPI2613968; A/American_crow/New_York/23-009035-001-original: EPI2613963, EPI2613964, EPI2613962, EPI2613966, EPI2613959, EPI2613965, EPI2613961, EPI2613960; A/red-tailed_hawk/Connecticut/23-009886-001-original: EPI2614123, EPI2614124, EPI2614122, EPI2614126, EPI2614119, EPI2614125, EPI2614121, EPI2614120; A/red-tailed_hawk/Connecticut/23-009880-001-original: EPI2614115, EPI2614116, EPI2614114, EPI2614118, EPI2614111, EPI2614117, EPI2614113, EPI2614112; A/turkey_vulture/New_York/23-011005-001-original: EPI2614171, EPI2614172, EPI2614170, EPI2614174, EPI2614167, EPI2614173, EPI2614169, EPI2614168; A/red-shouldered_hawk/New_York/23-012563-001-original: EPI2614339, EPI2614340, EPI2614338, EPI2614342, EPI2614335, EPI2614341, EPI2614337, EPI2614336; A/Canada_goose/New_York/23-013244-001-original: EPI2614395, EPI2614396, EPI2614394, EPI2614398, EPI2614391, EPI2614397, EPI2614393, EPI2614392; A/American_crow/New_York/23-014010-001-original: EPI2614427, EPI2614428, EPI2614426, EPI2614430, EPI2614423, EPI2614429, EPI2614425, EPI2614424; A/Canada_goose/New_York/23-014011-001-original: EPI2614435, EPI2614436, EPI2614434, EPI2614438, EPI2614431, EPI2614437, EPI2614433, EPI2614432

## Acknowledgments

We thank Wendy Puryear and Jonathan Runstadler from Tufts University for advice with the screening primers and protocols and staff at the Wild Bird Fund – specially Emily Einhorn and Rachel Frank - and the Animal Care Centers of New York City (Robin Brennen and Kevin Sexton) for providing samples, data and sampling expertise. We also would like to thank Dr. Sally Slavinsky from the New York City Department of Health and Mental Hygiene for advise and input, the NYC Department of Education, the NYC Parks and Recreation for sampling permits, Mary Pantin-Jackwood from USDA ARS for advice on genotyping, Isabel Francisco for her work on the initial two years of the New York Virus Hunters Program, Viviana Simon with help regarding regulatory questions and anonymous bird rescues. We would like to thank Flu Lab for funding this program and Julie Schafer and Casey Wright for their enthusiastic support. Furthermore, we would like to thank anonymous donors and Peter Palese for donations that made our annual New York City Virus Hunters Symposium possible. In addition, we would like to thank the Richard Lounsbery Foundation for project support. The enhanced Emerging Pathogens Facility (EPF) utilized in this project to deposit positive samples is a NIHBSL3/BSL3/ABSL3 facility that is part of the BSL-3 Biocontainment CoRE. This core is supported by subsidies from the ISMMS Dean’s Office and by investigator support through a cost recovery mechanism. Facility use reported in this publication was supported by the National Institute of Allergy And Infectious Diseases of the National Institutes of Health under Award Number G20AI174733 (R.A. Albrecht). The content is solely the responsibility of the authors and does not necessarily represent the official views of the National Institutes of Health. We gratefully acknowledge all data contributors, i.e., the Authors and their Originating laboratories responsible for obtaining the specimens, and their Submitting laboratories for generating the genetic sequence and metadata and sharing via the GISAID Initiative, which were used for comparisons with our sequences.

## Data availability statement

Sequences have been uploaded to GenBank and can be retrieved under the following identifiers: OR818561-OR81856, OR818637 - OR818644, OR818684 - OR818691, OR819057-OR819064, OR858836 - OR858843, OR819337 - OR819344.

## Figure legends

**Supplemental Figure 1.**
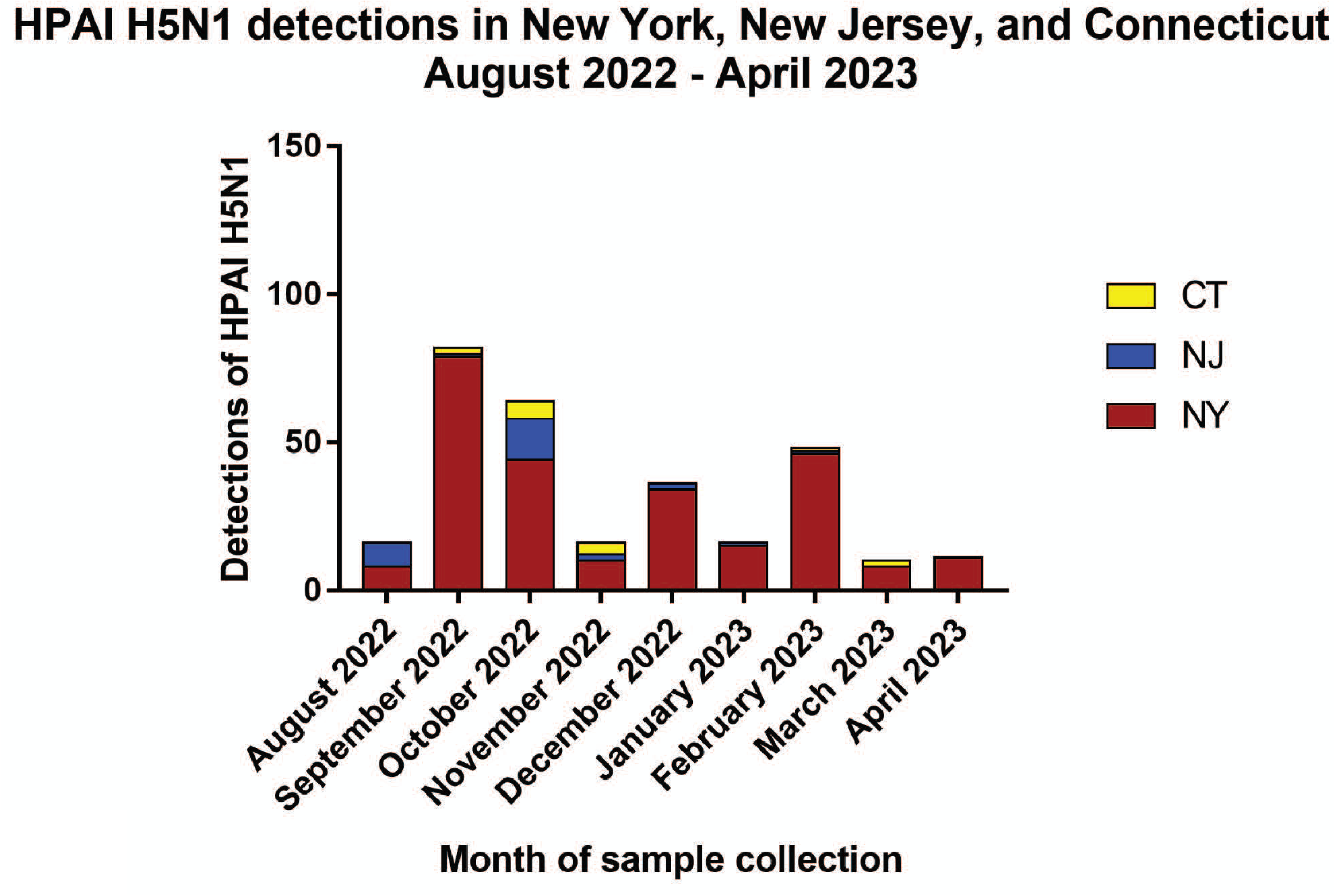
HPAI H5N1 detections in New York, New Jersey, and Connecticut, August 2022 – April 2023. Detections of HPAI H5N1 from the area surrounding New York City during the time period in which HPAI H5N1 was detected in New York City birds by NYCVH. Data retrieved on from the USDA APHIS database on wild bird HPAI H5N1 detections.

## References

1. Claas EC, Osterhaus AD, van Beek R, De Jong JC, Rimmelzwaan GF, Senne DA, Krauss S, Shortridge KF, Webster RG. 1998. Human influenza A H5N1 virus related to a highly pathogenic avian influenza virus. Lancet 351:472–7.

2. Xu X, Subbarao, Cox NJ, Guo Y. 1999. Genetic characterization of the pathogenic influenza A/Goose/Guangdong/1/96 (H5N1) virus: similarity of its hemagglutinin gene to those of H5N1 viruses from the 1997 outbreaks in Hong Kong. Virology 261:15–9.

3. (CDC) CfDCaP. 2004. Outbreaks of avian influenza A (H5N1) in Asia and interim recommendations for evaluation and reporting of suspected cases--United States, 2004. MMWR Morb Mortal Wkly Rep 53:97–100.

4. Abdelwhab EM, Selim AA, Arafa A, Galal S, Kilany WH, Hassan MK, Aly MM, Hafez MH. 2010. Circulation of avian influenza H5N1 in live bird markets in Egypt. Avian Dis 54:911–4.

5. Kayali G, Webby RJ, Ducatez MF, El Shesheny RA, Kandeil AM, Govorkova EA, Mostafa A, Ali MA. 2011. The epidemiological and molecular aspects of influenza H5N1 viruses at the human-animal interface in Egypt. PLoS One 6:e17730.

6. Terregino C, Milani A, Capua I, Marino AM, Cavaliere N. 2006. Highly pathogenic avian influenza H5N1 subtype in mute swans in Italy. Vet Rec 158:491.

7. Mølbak K, Trykker H, Mellergaard S, Glismann S. 2006. Avian influenza in Denmark, March-June 2006: public health aspects. Euro Surveill 11:E060615.3.

8. Enserink M. 2006. Avian influenza. H5N1 moves into Africa, European Union, deepening global crisis. Science 311:932.

9. Ducatez MF, Olinger CM, Owoade AA, De Landtsheer S, Ammerlaan W, Niesters HG, Osterhaus AD, Fouchier RA, Muller CP. 2006. Avian flu: multiple introductions of H5N1 in Nigeria. Nature 442:37.

10. Claes F, Morzaria SP, Donis RO. 2016. Emergence and dissemination of clade 2.3.4.4 H5Nx influenza viruses-how is the Asian HPAI H5 lineage maintained. Curr Opin Virol 16:158–163.

11. Krauss S, Stallknecht DE, Slemons RD, Bowman AS, Poulson RL, Nolting JM, Knowles JP, Webster RG. 2016. The enigma of the apparent disappearance of Eurasian highly pathogenic H5 clade 2.3.4.4 influenza A viruses in North American waterfowl. Proc Natl Acad Sci U S A 113:9033–8.

12. Adlhoch C, Fusaro A, Gonzales JL, Kuiken T, Marangon S, Niqueux É, Staubach C, Terregino C, Aznar I, Muñoz Guajardo I, Baldinelli F, European Food Safety Authority ECfDP, Control, E.ropean Union Reference Laboratory for Avian Influenza. 2021. Avian influenza overview September - December 2021. EFSA J 19:e07108.

13. Caliendo V, Lewis NS, Pohlmann A, Baillie SR, Banyard AC, Beer M, Brown IH, Fouchier RAM, Hansen RDE, Lameris TK, Lang AS, Laurendeau S, Lung O, Robertson G, van der Jeugd H, Alkie TN, Thorup K, van Toor ML, Waldenström J, Yason C, Kuiken T, Berhane Y. 2022. Transatlantic spread of highly pathogenic avian influenza H5N1 by wild birds from Europe to North America in 2021. Sci Rep 12:11729.

14. Günther A, Krone O, Svansson V, Pohlmann A, King J, Hallgrimsson GT, Skarphéðinsson KH, Sigurðardóttir H, Jónsson SR, Beer M, Brugger B, Harder T. 2022. Iceland as Stepping Stone for Spread of Highly Pathogenic Avian Influenza Virus between Europe and North America. Emerg Infect Dis 28:2383–2388.

15. Alkie TN, Lopes S, Hisanaga T, Xu W, Suderman M, Koziuk J, Fisher M, Redford T, Lung O, Joseph T, Himsworth CG, Brown IH, Bowes V, Lewis NS, Berhane Y. 2022. A threat from both sides: Multiple introductions of genetically distinct H5 HPAI viruses into Canada via both East Asia-Australasia/Pacific and Atlantic flyways. Virus Evol 8:veac077.

16. Youk S, Torchetti MK, Lantz K, Lenoch JB, Killian ML, Leyson C, Bevins SN, Dilione K, Ip HS, Stallknecht DE, Poulson RL, Suarez DL, Swayne DE, Pantin-Jackwood MJ. 2023. H5N1 highly pathogenic avian influenza clade 2.3.4.4b in wild and domestic birds: Introductions into the United States and reassortments, December 2021-April 2022. Virology 587:109860.

17. Duriez O, Sassi Y, Le Gall-Ladevèze C, Giraud L, Straughan R, Dauverné L, Terras A, Boulinier T, Choquet R, Van De Wiele A, Hirschinger J, Guérin JL, Le Loc’h G. 2023. Highly pathogenic avian influenza affects vultures’ movements and breeding output. Curr Biol 33:3766–3774.e3.

18. Rijks JM, Leopold MF, Kühn S, In ’t Veld R, Schenk F, Brenninkmeijer A, Lilipaly SJ, Ballmann MZ, Kelder L, de Jong JW, Courtens W, Slaterus R, Kleyheeg E, Vreman S, Kik MJL, Gröne A, Fouchier RAM, Engelsma M, de Jong MCM, Kuiken T, Beerens N. 2022. Mass Mortality Caused by Highly Pathogenic Influenza A(H5N1) Virus in Sandwich Terns, the Netherlands, 2022. Emerg Infect Dis 28:2538–2542.

19. Gamarra-Toledo V, Plaza PI, Angulo F, Gutiérrez R, García-Tello O, Saravia-Guevara P, Mejía-Vargas F, Epiquién-Rivera M, Quiroz-Jiménez G, Martinez P, Huamán-Mendoza D, Inga-Díaz G, La Madrid LE, Luyo P, Ventura S, Lambertucci SA. 2023. Highly Pathogenic Avian Influenza (HPAI) strongly impacts wild birds in Peru. Biological Conservation 286:110272.

20. Jakobek BT, Berhane Y, Nadeau MS, Embury-Hyatt C, Lung O, Xu W, Lair S. 2023. Influenza A(H5N1) Virus Infections in 2 Free-Ranging Black Bears (Ursus americanus), Quebec, Canada. Emerg Infect Dis 29:2145–2149.

21. Nguyen HT, Chesnokov A, De La Cruz J, Pascua PNQ, Mishin VP, Jang Y, Jones J, Di H, Ivashchenko AA, Killian ML, Torchetti MK, Lantz K, Wentworth DE, Davis CT, Ivachtchenko AV, Gubareva LV. 2023. Antiviral susceptibility of clade 2.3.4.4b highly pathogenic avian influenza A(H5N1) viruses isolated from birds and mammals in the United States, 2022. Antiviral Res 217:105679.

22. Alkie TN, Cox S, Embury-Hyatt C, Stevens B, Pople N, Pybus MJ, Xu W, Hisanaga T, Suderman M, Koziuk J, Kruczkiewicz P, Nguyen HH, Fisher M, Lung O, Erdelyan CNG, Hochman O, Ojkic D, Yason C, Bravo-Araya M, Bourque L, Bollinger TK, Soos C, Giacinti J, Provencher J, Ogilvie S, Clark A, MacPhee R, Parsons GJ, Eaglesome H, Gilbert S, Saboraki K, Davis R, Jerao A, Ginn M, Jones MEB, Berhane Y. 2023. Characterization of neurotropic HPAI H5N1 viruses with novel genome constellations and mammalian adaptive mutations in free-living mesocarnivores in Canada. Emerg Microbes Infect 12:2186608.

23. Leguia M, Garcia-Glaessner A, Muñoz-Saavedra B, Juarez D, Barrera P, Calvo-Mac C, Jara J, Silva W, Ploog K, Amaro L, Colchao-Claux P, Johnson CK, Uhart MM, Nelson MI, Lescano J. 2023. Highly pathogenic avian influenza A (H5N1) in marine mammals and seabirds in Peru. Nat Commun 14:5489.

24. Ulloa M, Fernández A, Ariyama N, Colom-Rivero A, Rivera C, Nuñez P, Sanhueza P, Johow M, Araya H, Torres JC, Gomez P, Muñoz G, Agüero B, Alegría R, Medina R, Neira V, Sierra E. 2023. Mass mortality event in South American sea lions (Vet Q 43:1–10.

25. Puryear W, Sawatzki K, Hill N, Foss A, Stone JJ, Doughty L, Walk D, Gilbert K, Murray M, Cox E, Patel P, Mertz Z, Ellis S, Taylor J, Fauquier D, Smith A, DiGiovanni RA, van de Guchte A, Gonzalez-Reiche AS, Khalil Z, van Bakel H, Torchetti MK, Lantz K, Lenoch JB, Runstadler J. 2023. Highly Pathogenic Avian Influenza A(H5N1) Virus Outbreak in New England Seals, United States. Emerg Infect Dis 29:786–791.

26. Lindh E, Lounela H, Ikonen N, Kantala T, Savolainen-Kopra C, Kauppinen A, Österlund P, Kareinen L, Katz A, Nokireki T, Jalava J, London L, Pitkäpaasi M, Vuolle J, Punto-Luoma AL, Kaarto R, Voutilainen L, Holopainen R, Kalin-Mänttäri L, Laaksonen T, Kiviranta H, Pennanen A, Helve O, Laamanen I, Melin M, Tammiranta N, Rimhanen-Finne R, Gadd T, Salminen M. 2023. Highly pathogenic avian influenza A(H5N1) virus infection on multiple fur farms in the South and Central Ostrobothnia regions of Finland, July 2023. Euro Surveill 28.

27. Agüero M, Monne I, Sánchez A, Zecchin B, Fusaro A, Ruano MJ, Del Valle Arrojo M, Fernández-Antonio R, Souto AM, Tordable P, Cañás J, Bonfante F, Giussani E, Terregino C, Orejas JJ. 2023. Highly pathogenic avian influenza A(H5N1) virus infection in farmed minks, Spain, October 2022. Euro Surveill 28.

28. Maemura T, Guan L, Gu C, Eisfeld A, Biswas A, Halfmann P, Neumann G, Kawaoka Y. 2023. Characterization of highly pathogenic clade 2.3.4.4b H5N1 mink influenza viruses. EBioMedicine 97:104827.

29. Castillo A, Fasce R, Parra B, Andrade W, Covarrubias P, Hueche A, Campano C, Tambley C, Rojas M, Araya M, Hernández F, Bustos P, Fernández J. 2023. The first case of human infection with H5N1 avian Influenza A virus in Chile. J Travel Med 30.

30. Bruno A, Alfaro-Núñez A, de Mora D, Armas R, Olmedo M, Garcés J, Garcia-Bereguiain MA. 2023. First case of human infection with highly pathogenic H5 avian Influenza A virus in South America: A new zoonotic pandemic threat for 2023? J Travel Med 30.

31. CDC. 2023. https://www.cdc.gov/flu/avianflu/spotlights/2022-2023/h5n1-technical-report_october.htm.

32. Sillman SJ, Drozd M, Loy D, Harris SP. 2023. Naturally occurring highly pathogenic avian influenza virus H5N1 clade 2.3.4.4b infection in three domestic cats in North America during 2023. J Comp Pathol 205:17–23.

33. Domańska-Blicharz K, Świętoń E, Świątalska A, Monne I, Fusaro A, Tarasiuk K, Wyrostek K, Styś-Fijoł N, Giza A, Pietruk M, Zechchin B, Pastori A, Adaszek Ł, Pomorska-Mól M, Tomczyk G, Terregino C, Winiarczyk S. 2023. Outbreak of highly pathogenic avian influenza A(H5N1) clade 2.3.4.4b virus in cats, Poland, June to July 2023. Euro Surveill 28.

34. Briand FX, Souchaud F, Pierre I, Beven V, Hirchaud E, Hérault F, Planel R, Rigaudeau A, Bernard-Stoecklin S, Van der Werf S, Lina B, Gerbier G, Eterradossi N, Schmitz A, Niqueux E, Grasland B. 2023. Highly Pathogenic Avian Influenza A(H5N1) Clade 2.3.4.4b Virus in Domestic Cat, France, 2022. Emerg Infect Dis 29:1696–1698.

35. Moreno A, Bonfante F, Bortolami A, Cassaniti I, Caruana A, Cottini V, Cereda D, Farioli M, Fusaro A, Lavazza A, Lecchini P, Lelli D, Maroni Ponti A, Nassuato C, Pastori A, Rovida F, Ruocco L, Sordilli M, Baldanti F, Terregino C. 2023. Asymptomatic infection with clade 2.3.4.4b highly pathogenic avian influenza A(H5N1) in carnivore pets, Italy, April 2023. Euro Surveill 28.

36. Francisco I, Bailey S, Bautista T, Diallo D, Gonzalez J, Kirkpatrick Roubidoux E, Kehinde Ajayi P, Albrecht RA, McMahon R, Krammer F, Marizzi C. 2022. Detection of Velogenic Avian Paramyxoviruses in Rock Doves in New York City, New York. Microbiol Spectr 10:e0206121.

37. Dickinson JL, Shirk J, Bonter D, Bonney R, Crain RL, Martin J, Phillips T, Purcell K. 2012. The current state of citizen science as a tool for ecological research and public engagement. Frontiers in Ecology and the Environment 10:291–297.

38. Saavedra I, Rabadán-González J, Aragonés D, Figuerola J. 2023. Can Citizen Science Contribute to Avian Influenza Surveillance? Pathogens 12.

39. Tan YR, Agrawal A, Matsoso MP, Katz R, Davis SLM, Winkler AS, Huber A, Joshi A, El-Mohandes A, Mellado B, Mubaira CA, Canlas FC, Asiki G, Khosa H, Lazarus JV, Choisy M, Recamonde-Mendoza M, Keiser O, Okwen P, English R, Stinckwich S, Kiwuwa-Muyingo S, Kutadza T, Sethi T, Mathaha T, Nguyen VK, Gill A, Yap P. 2022. A call for citizen science in pandemic preparedness and response: beyond data collection. BMJ Glob Health 7.

40. Marizzi C, Wright L. 2024. Hunting emerging viruses through participatory community science. Nat Microbiol 9:578–581.

41. Kandeil A, Patton C, Jones JC, Jeevan T, Harrington WN, Trifkovic S, Seiler JP, Fabrizio T, Woodard K, Turner JC, Crumpton JC, Miller L, Rubrum A, DeBeauchamp J, Russell CJ, Govorkova EA, Vogel P, Kim-Torchetti M, Berhane Y, Stallknecht D, Poulson R, Kercher L, Webby RJ. 2023. Rapid evolution of A(H5N1) influenza viruses after intercontinental spread to North America. Nat Commun 14:3082.

42. Antanasijevic A, Durst MA, Cheng H, Gaisina IN, Perez JT, Manicassamy B, Rong L, Lavie A, Caffrey M. 2020. Structure of avian influenza hemagglutinin in complex with a small molecule entry inhibitor. Life Sci Alliance 3.

43. Cattoli G, Milani A, Temperton N, Zecchin B, Buratin A, Molesti E, Aly MM, Arafa A, Capua I. 2011. Antigenic drift in H5N1 avian influenza virus in poultry is driven by mutations in major antigenic sites of the hemagglutinin molecule analogous to those for human influenza virus. J Virol 85:8718–24.

44. Ohshima N, Iba Y, Kubota-Koketsu R, Yamasaki A, Majima K, Kurosawa G, Hirano D, Yoshida S, Sugiura M, Asano Y, Okuno Y, Kurosawa Y. 2020. Comprehensive Analysis of Antibodies Induced by Vaccination with 4 Kinds of Avian Influenza H5N1 Pre-Pandemic Vaccines. Int J Mol Sci 21.

45. CDC. Accessed on 4/2/24. https://www.cdc.gov/flu/avianflu/communication-resources/index.html.

46. FAO. Accessed on 4/2/24. https://www.cms.int/sites/default/files/publication/avian_influenza_2023_aug.pdf.

47. Questa K, Das M, King R, Everitt M, Rassi C, Cartwright C, Ferdous T, Barua D, Putnis N, Snell AC, Huque R, Newell J, Elsey H. 2020. Community engagement interventions for communicable disease control in low- and lower-middle-income countries: evidence from a review of systematic reviews. Int J Equity Health 19:51.

48. Krammer F, Schultz-Cherry S. 2023. We need to keep an eye on avian influenza. Nat Rev Immunol 23:267–268.

49. Xie R, Edwards KM, Wille M, Wei X, Wong SS, Zanin M, El-Shesheny R, Ducatez M, Poon LLM, Kayali G, Webby RJ, Dhanasekaran V. 2023. The episodic resurgence of highly pathogenic avian influenza H5 virus. Nature 622:810–817.

50. Institute TA. 2023. https://www.aspeninstitute.org/wp-content/uploads/2023/12/WHY-and-HOW-of-Public-Trust-in-Science-Aspen-Institute.pdf.

51. Stoto MA, Schlageter S, Kraemer JD. 2022. COVID-19 mortality in the United States: It’s been two Americas from the start. PLoS One 17:e0265053.

52. Pasick J, Berhane Y, Embury-Hyatt C, Copps J, Kehler H, Handel K, Babiuk S, Hooper-McGrevy K, Li Y, Mai Le Q, Lien Phuong S. 2007. Susceptibility of Canada Geese (Branta canadensis) to highly pathogenic avian influenza virus (H5N1). Emerg Infect Dis 13:1821–7.

53. Neufeld JL, Embury-Hyatt C, Berhane Y, Manning L, Ganske S, Pasick J. 2009. Pathology of highly pathogenic avian influenza virus (H5N1) infection in Canada geese (Branta canadensis): preliminary studies. Vet Pathol 46:966–70.

54. Guardian T. 2023. Pet dog dies after catching bird flu, vol https://www.telegraph.co.uk/global-health/science-and-disease/bird-flu-cases-pet-dog-canada-vaccine/.

55. Hoffmann E, Stech J, Guan Y, Webster RG, Perez DR. 2001. Universal primer set for the full-length amplification of all influenza A viruses. Arch Virol 146:2275–89.

56. Kim GS, Kim TS, Son JS, Lai VD, Park JE, Wang SJ, Jheong WH, Mo IP. 2019. The difference of detection rate of avian influenza virus in the wild bird surveillance using various methods. J Vet Sci 20:e56.

57. WHO. 2021. WHO information for the molecular detection of influenza viruses, vol https://cdn.who.int/media/docs/default-source/influenza/molecular-detention-of-influenza-viruses/protocols_influenza_virus_detection_feb_2021.pdf.

58. Afgan E, Baker D, van den Beek M, Blankenberg D, Bouvier D, Čech M, Chilton J, Clements D, Coraor N, Eberhard C, Grüning B, Guerler A, Hillman-Jackson J, Von Kuster G, Rasche E, Soranzo N, Turaga N, Taylor J, Nekrutenko A, Goecks J. 2016. The Galaxy platform for accessible, reproducible and collaborative biomedical analyses: 2016 update. Nucleic Acids Res 44:W3–W10.

59. Prjibelski A, Antipov D, Meleshko D, Lapidus A, Korobeynikov A. 2020. Using SPAdes De Novo Assembler. Curr Protoc Bioinformatics 70:e102.

60. Bao Y, Bolotov P, Dernovoy D, Kiryutin B, Zaslavsky L, Tatusova T, Ostell J, Lipman D. 2008. The influenza virus resource at the National Center for Biotechnology Information. J Virol 82:596–601.

61. Vasimuddin M, Misra S, Li H, Aluru S. 2019. Efficient Architecture-Aware Acceleration of BWA-MEM for Multicore Systems, abstr 2019 IEEE International Parallel and Distributed Processing Symposium (IPDPS), IEEE Computer Society,

62. Langmead B, Salzberg SL. 2012. Fast gapped-read alignment with Bowtie 2. Nat Methods 9:357–9.

63. Grubaugh ND, Gangavarapu K, Quick J, Matteson NL, De Jesus JG, Main BJ, Tan AL, Paul LM, Brackney DE, Grewal S, Gurfield N, Van Rompay KKA, Isern S, Michael SF, Coffey LL, Loman NJ, Andersen KG. 2019. An amplicon-based sequencing framework for accurately measuring intrahost virus diversity using PrimalSeq and iVar. Genome Biol 20:8.

64. Edgar RC. 2004. MUSCLE: multiple sequence alignment with high accuracy and high throughput. Nucleic Acids Res 32:1792–7.

65. Tamura K, Nei M. 1993. Estimation of the number of nucleotide substitutions in the control region of mitochondrial DNA in humans and chimpanzees. Mol Biol Evol 10:512–26.

